# Parallelized engineering of mutational models using piggyBac transposon delivery of CRISPR libraries

**DOI:** 10.1101/2020.07.10.197962

**Authors:** Xander Nuttle, Nicholas D. Burt, Benjamin Currall, Mariana Moysés-Oliveira, Kiana Mohajeri, Rachita Yadav, Derek J. C. Tai, James F. Gusella, Michael E. Talkowski

## Abstract

Novel gene and variant discoveries have reached unprecedented scale with the emergence of exome and genome sequencing studies across a spectrum of human disease initiatives. Highly parallelized functional characterization of these variants is now paramount to deciphering disease mechanisms, and approaches that facilitate editing of induced pluripotent stem cells (iPSCs) to derive otherwise inaccessible tissues of interest (e.g., brain) have become critical in genomics research. Here, we sought to facilitate scalable editing of multiple genes and variants by developing a genome engineering approach that incorporates libraries of CRISPR/Cas9 guide RNAs (gRNAs) into a piggyBac (PB) transposon system. To test the efficiency of inducing small indels, targeted deletions, and large reciprocal copy number variants (CNVs), we simultaneously delivered to human iPSCs both *Cas9* and a library including 59 single gRNAs targeting segmental duplications, 70 paired gRNAs flanking particular genic regions, and three single gRNAs targeting the coding sequence of an individual gene, *MAGEL2*. After editing, we isolated single cells, expanded resultant colonies, and genotyped their gRNA contents and mutational outcomes. We observed that 97.7% of gRNA constructs were integrated into at least one colony, with 85.6% of colonies containing three or fewer PB integrations. This PB editing method generated 354 cell lines with 57.8% of sequenced gRNA cleavage sites modified in at least one line, 14.4% of these lines altered at multiple targets, and single-copy indel mutagenesis predominating. Among the edits generated were eight targeted genomic deletions, including pathogenic microdeletions at chromosome 15q11-q13 (∼5.3 Mbp), chromosome 16p11.2 (∼740 kbp), and chromosome 17q11.2 (∼1.4 Mbp). These data highlight PB editing as a powerful platform for gene inactivation and testify to its strong potential for oligogenic modeling. The ability to rapidly establish high-quality mutational models at scale will facilitate the development of near-isogenic cellular collections and catalyze comparative functional genomic studies, better positioning us to investigate the roles of hundreds of genes and mutations in development and disease.

---

A major challenge in human genomics is translating statistical associations of genetic variants into functional biological insights and pathogenic mechanisms^1^. The increasing scale of genome and exome sequencing studies has produced a deluge of gene and variant associations across Mendelian and complex disorders^2-5^, yet forging mechanistic links from genotypes to phenotypes remains daunting. Until recently, most mechanistic studies have relied upon patient-derived cell lines or model organisms^6,7^. These approaches can have limitations in the number of patients that can be recruited with the desired variants, access to clinically relevant tissues, and difficulties in interpretation arising from variable genetic backgrounds or uncertain generalizability of findings in animals for humans^8,9^. Consequently, there is a critical need for approaches that can rapidly model disease-associated variants and perturb functional networks to inform therapeutic targeting and development.

The emergence of iPSC and genome engineering technologies over the past decade offers unprecedented opportunities to experimentally interrogate mutations of interest^10,11^. An expanding compendium of *in vitro* differentiation protocols has enabled researchers to routinely cultivate a wide variety of cell types in a dish, including several distinct populations of neurons^12,13^. Three-dimensional cell culture systems have also yielded organoids that recapitulate some hallmark properties of their *in vivo* counterparts, including cerebral organoids as models of brain development^13,14^. Genome editing tools, most notably CRISPR/Cas9 and its relatives^15,16^, provide methods to engineer a growing catalog of precisely targeted mutations, including indels^17-19^, point mutations^20,21^, CNVs^22,23^, and inversions^23,24^. Continued technological developments such as prime editing promise to enhance editing efficiencies, augment targeting possibilities, and expand the range of programmable mutations^25,26^.

To fully capitalize on these advances for cellular mutational modeling, particularly for strategies involving genome engineering in iPSCs to establish clonal mutant lines, approaches to parallelize and scale the experimental workflow are needed. It is currently possible to generate multiple programmed edits by performing separate transfections with different gRNAs, but time, cost, and labor considerations preclude using this approach for generating hundreds of individual mutations. Saturation genome editing has yielded cell populations with 3,893 coding single-nucleotide variants (SNVs), including 96.5% of all possible SNVs within thirteen targeted BRCA1 exons^27,28^. Although this strategy is extremely powerful for introducing desired genetic changes at a single locus, modifying other sites, even within the same gene, has required separate gRNA transfections. Lentiviral gRNA libraries have been successfully employed in genome-scale genetic screens, including attempts to knock out 19,050 human genes and 20,611 mouse genes^29-32^. However, resulting edited cells harbor integrated lentiviral sequence in addition to any CRISPR-mediated mutations, complicating efforts to infer their functional consequences. Moreover, none of these large-scale editing strategies has yet been coupled with isolating individual mutant cells to create cellular models.

Here we leveraged the PB transposon, a mobile genetic element that can be scarlessly removed from the cellular genome^33^, to deliver *Cas9*^34-37^ together with a gRNA library to iPSCs and perform parallelized genome engineering. We explored both CNV and indel formation, balancing our interest in modeling reciprocal genomic disorders (RGDs) and dosage sensitivity with our desire to develop a method with broad applicability. We evaluated editing outcomes by using molecular inversion probes (MIPs)^38-40^ and massively parallel sequencing to genotype clonal lines expanded from single cells. The method efficiently produced 56 cellular models with indels as well as 8 models with genomic deletions in a single two-month experiment.

## RESULTS

### Genome engineering strategy

We postulated that the PB transposon platform is well-suited for engineering cellular models in parallel for three reasons. First, PB transposons can be scarlessly excised from cellular genomes^33^, such that final models would genetically differ only by CRISPR-induced mutations^34^. Second, the PB system has previously been utilized for genome engineering and more recently for CRISPR applications^35-37^, having demonstrated efficient editing in iPSCs with indel frequencies of 40-50%^34^. Third, adjusting the amounts of transposon and transposase plasmids transfected allows some control over how many PB integrations occur per cell^35,41^. Thus, numbers of gRNAs delivered to each cell can be limited to reduce the possibility of models acquiring multiple mutations or increased for oligogenic studies. Accordingly, we developed a PB genome engineering workflow that incorporates delivery of editing machinery to iPSCs and its later removal, temporally-controlled editing, and screening for mutations of interest (**Fig. 1**). Initially, we clone gRNAs into a plasmid-based PB transposon containing inducible *Cas9* and a puromycin resistance gene. Transfecting the resulting CRISPR library into iPSCs together with a plasmid encoding PB transposase yields a mixed population of cells, with several integrating one or more PB transposons and constitutively expressing one or more gRNAs. Puromycin treatment selects for PB-containing cells, and addition of doxycycline to the culture medium induces editing. We then transfect remaining iPSCs with a plasmid encoding green fluorescent protein (GFP) and excision-only PB transposase and perform fluorescence-activated cell sorting (FACS) to isolate single cells where PB integrations were likely removed. After expanding colonies, we split them into separate plates for DNA extraction and propagation. MIP genotyping at gRNA cleavage sites identifies indels, while data from MIPs targeting SNVs across regions of interest allow detection of CNVs based on altered allele balance^38-40^.

**Figure 1.**
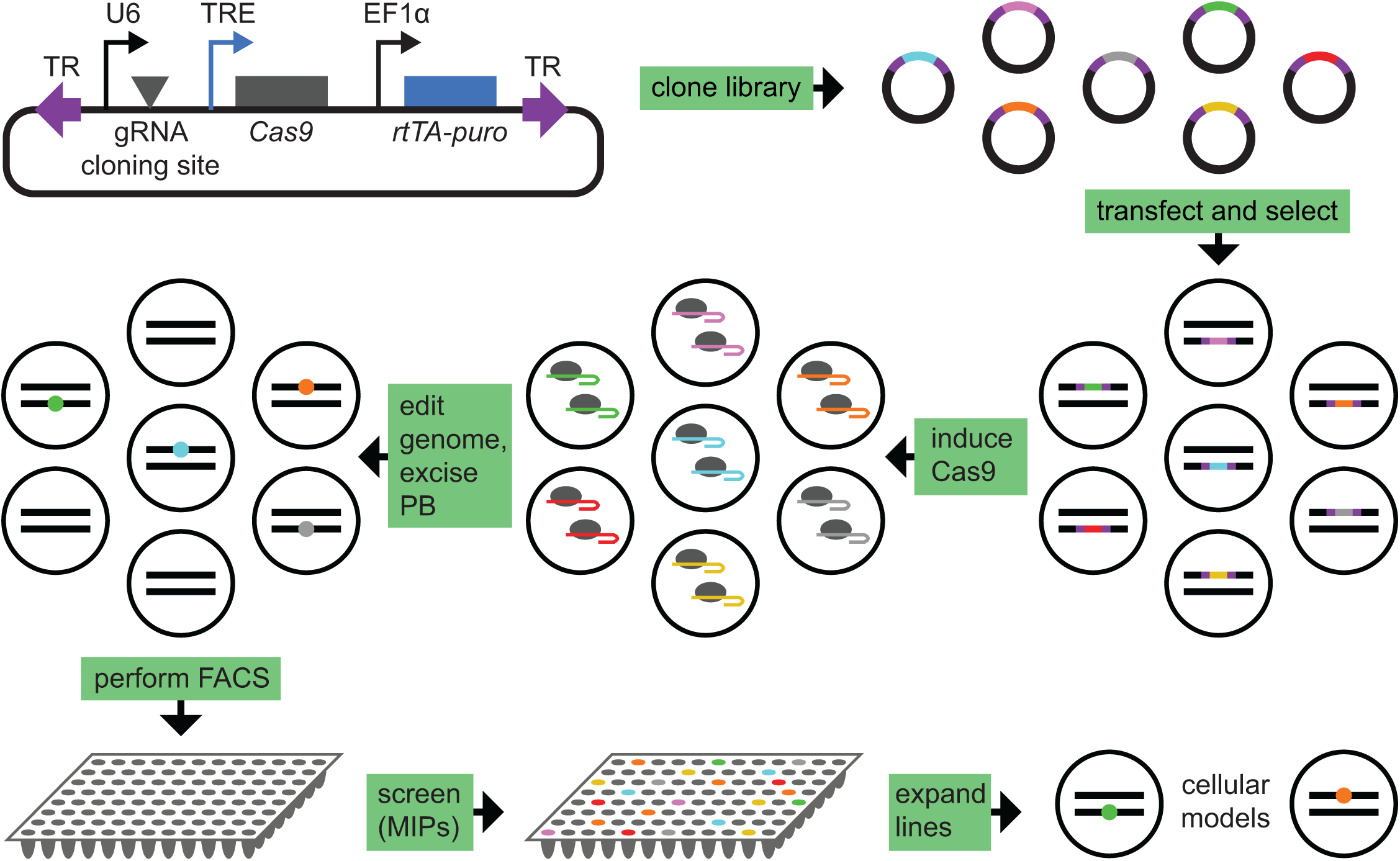
PB system for parallelized genome engineering. Schematic outlines experimental workflow. (Top) cloning a gRNA library into the PB plasmid vector. (Middle) delivering and removing CRISPR components via PB transposition enables multiplexed, temporally controlled editing. (Bottom) isolating single cells establishes clonal iPSC lines, which are rapidly genotyped using MIPs to identify mutational models. TR: terminal repeat; TRE: tetracycline response element; rtTA: reverse tetracycline-controlled transactivator.

### PB integration copy number and gRNA representation

To test this approach, we cloned a library that included 62 individual gRNAs and 70 gRNA pairs (**Supplementary Table 1**) into our PB vector and attempted editing as described above. In this library, all single gRNAs are driven by the U6 promotor in the vector. When the vector contains a pair of gRNAs, the first is driven by the U6 promotor while the second is driven by an inserted H1 promotor (see **Methods**). Nearly all individual gRNAs (59 of 62) targeted segmental duplications flanking known RGD loci to promote CNV formation^22^, while the remainder targeted *MAGEL2* coding sequence to generate loss-of-function indels. Most gRNA pairs (48 of 70) were designed for individual gene deletions, with others conducive to deleting multigene segments or the Prader-Willi syndrome imprinting center. We evaluated four library transfection conditions, lipofecting iPSCs with 125 ng, 250 ng, 500 ng, or 1000 ng of library plasmid DNA together with 1000 ng PB transposase plasmid. Corresponding control transfections included 1000 ng PB library DNA or 1000 ng PB transposase plasmid. After three days of puromycin treatment, all colonies had detached from control wells, while all other conditions yielded large numbers of resistant colonies. We omitted PB excision and genotyped gRNA contents for each expanded colony, enabling estimation of PB integration copy numbers and comparisons of potential edits with observed mutations. Interestingly, different DNA inputs showed no significant differences in PB integration copy numbers per cell (**Fig. 2a**; k-sample Anderson-Darling p = 0.2, 1,000,000 simulations), estimated by counting distinct integrated gRNA constructs identified in clonal lines established from the transfected iPSCs without PB excision. PB transfections delivered 0-9 gRNA constructs to each cell, with 85.6% of lines with high-confidence genotypes (285 of 333) having received three or fewer integrations. Lines lacking PB integrations may have originated from cells protected from puromycin by nearby resistant colonies. Importantly, 90.7% of lines (321 of 354) showed no evidence of random plasmid integration based on detection of plasmid sequences within and outside the PB transposon (**Supplementary Fig. 1**).

**Figure 2.**
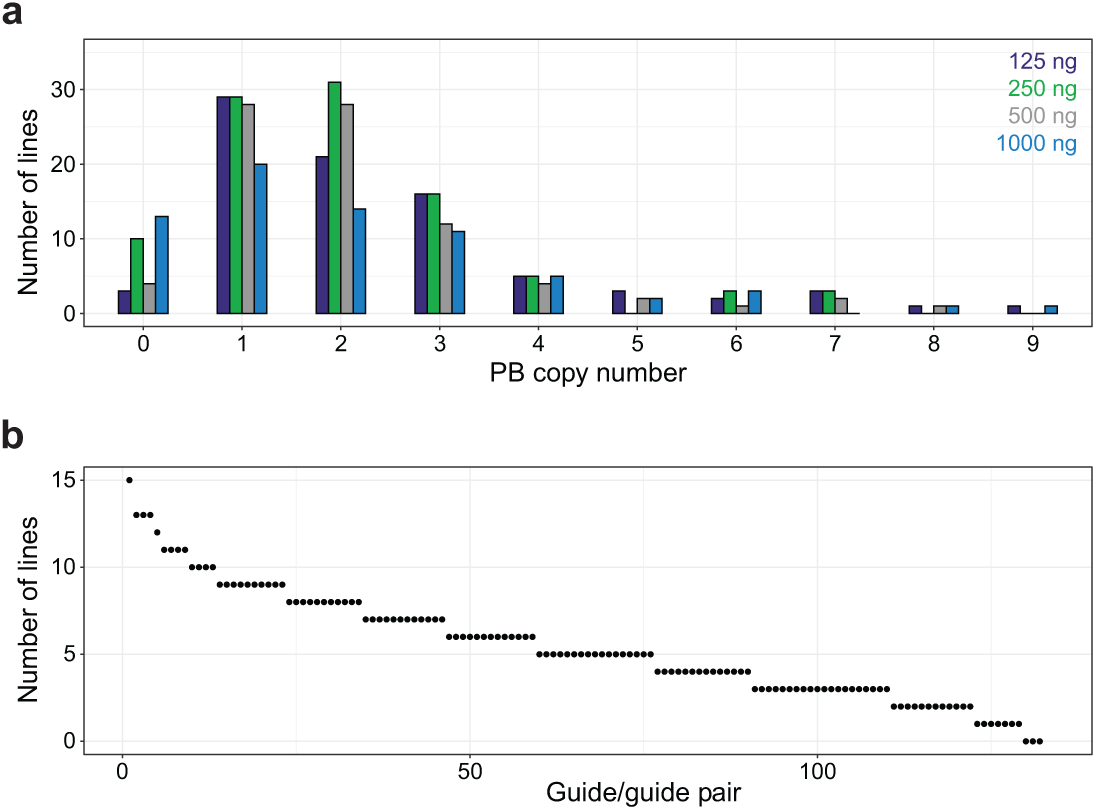
Characterization of PB integrations. a) Distributions of PB copy numbers per cell associated with four transfection conditions differing by amounts of PB gRNA plasmid library included. PB copy number was inferred by counting distinct integrated gRNA constructs. b) gRNA library representation. Points indicate how many iPSC lines contained each construct. Constructs are ranked by abundance. As expected, no line harbored an empty construct with stuffer sequence that was present prior to library cloning.

The likelihood of engineering a specific mutation of interest using our system depends in part on the relative integration frequency of the corresponding gRNA construct. For example, an inefficient gRNA might generate more mutant lines than a highly active gRNA if the former were expressed in more cells than the latter. Thus, to assess gRNA representation, we counted the number of isolated iPSC lines containing each gRNA construct. Per-construct integration-line counts ranged from 0-15 (mean = 5.47; median = 5.0; **Fig. 2b**), and their distribution was relatively consistent but statistically non-uniform (exact multinomial goodness-of-fit p < 10^−6^, 1,000,000 simulations; **Supplementary Fig. 2** and **Supplementary Tables 2-3**). From these analyses, 97.7% of constructs (129 of 132) were incorporated into at least one line, with only three constructs never detected, suggesting efficient delivery and good representation of the gRNA library.

### Generation of indels

Our genotyping assay included 43 MIPs that successfully captured 45 gRNA cleavage sites, allowing us to explore the performance of our method for multiplexed indel mutagenesis. Most of these cleavage sites (37) were in segmental duplications flanking RGD regions. Sequence analysis revealed that 57.8% of successfully captured gRNA targets (26 of 45) had inserted and/or deleted bases in at least one iPSC line (**Fig. 3a**). Considering only lines containing integrated gRNA constructs targeting sites captured by our MIP panel, 37.1% (56 of 151) were edited, with most of these (85.7%, 48 of 56) acquiring indels from activity of only one gRNA (**Fig. 3b**). Given that several gRNAs mediated efficient indel formation (**Supplementary Table 4**), we investigated whether editing more often altered single alleles or multiple target copies. Counting numbers of distinct indel-containing alignments associated with each target sequence in each edited iPSC line indicated that single-copy mutagenesis predominated (38 of 66 genotypes), although two- and three-copy editing occurred multiple times (**Supplementary Fig. 3**). Because most of these genotypes (59 of 66) involve multicopy duplicated sequences flanking RGD regions, observing more than one indel alignment likely reflects mutagenesis at distinct copies/sites rather than mosaicism from two or more indels at the same site. Notably, all editing in unique sequence resulted in single indels, with six apparently heterozygous clones showing indel-to-unedited allele balances between 0.45 and 0.57 and one likely mosaic (same allele balance 0.11). No indels resulted from gRNAs targeting *MAGEL2* coding sequence, but only two of their cleavage sites could be genotyped, and together, corresponding constructs were integrated in just three lines. We did not discern a noticeable trend between indel editing efficiencies and integrated PB copy numbers (**Supplementary Fig. 4**), although no analyzed lines had PB copy numbers above nine. In aggregate, we simultaneously generated 56 mutant iPSC lines harboring indels localized to 26 targets screened. An additional 224 iPSC lines together likely contain dozens of undetected indels since their PB integrations included at least one of 157 gRNA constructs targeting sites not captured by our MIP panel. These results highlight the power of PB genome engineering for high-throughput production of cellular models featuring indels, including oligogenic models and heterozygous models for haploinsufficiency studies.

**Figure 3.**
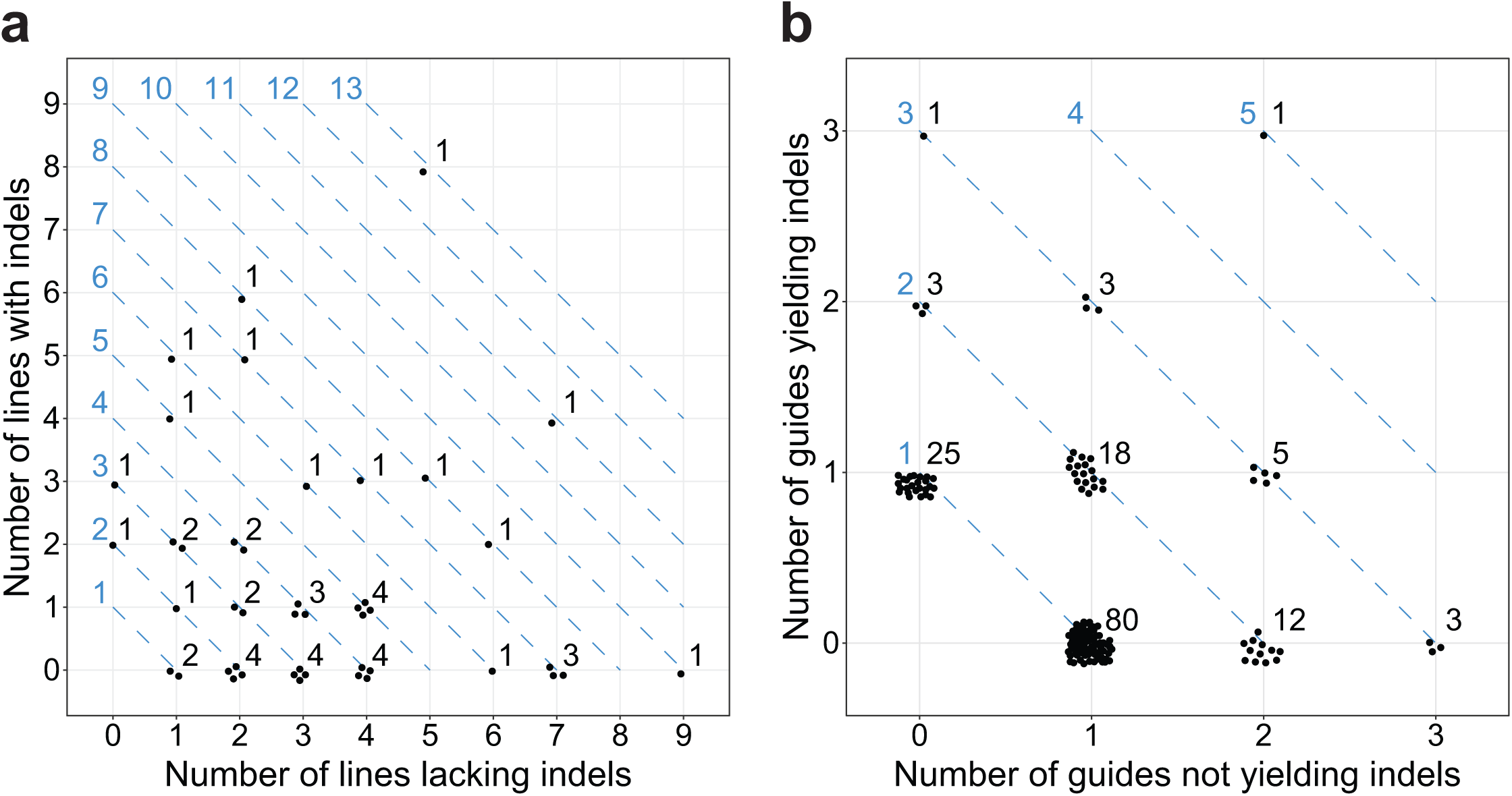
Summary of PB CRISPR indel mutagenesis. a) gRNAs (points) are grouped along diagonals (blue lines) based on numbers of lines expressing them (blue). Positions along each diagonal (jittered around their integer values) reflect how many such lines acquired corresponding indels. Black numbers quantify points at each position. b) iPSC lines (points) are grouped along diagonals (blue lines) based on numbers of gRNAs they express (blue) amenable to indel analysis. Positions along each diagonal (jittered around their integer values) indicate how many such gRNAs induced indels during editing with Cas9. Black numbers quantify points at each position.

### Production of multiple CNV microdeletions

Editing with single gRNAs targeting duplicated sequences or gRNA pairs occasionally results in two double-strand DNA breaks on a single chromosome, with imperfect repair sometimes leading to deletion or duplication of intervening sequence. We and others have previously leveraged these strategies to generate cellular models with CNVs equivalent to those found in RGD patients^22^ or single-gene deletions^23^, although never in a parallelized manner. PB editing yielded eight microdeletions of varying sizes (**Supplementary Table 5**), including programmed CNV microdeletions of chromosome 15q11-q13 (∼5.3 Mbp), chromosome 16p11.2 (∼740 kbp), and chromosome 17q11.2 (∼1.4 Mbp, **Fig. 4a**). The gRNA pairs yielded a single smaller deletion spanning *MAGEL2* (∼6.9 kbp; **Fig. 4b**). Finally, we observed two novel deletions resulting from CRISPR activity involving two distinct integrated gRNA constructs that happened to target the same chromosome, an ∼3.7 Mbp deletion at chromosome 15q11-q13 and an ∼214 kbp deletion at chromosome 16p11.2 (**Fig. 4c**). No duplications were identified, reflecting the expected lower efficiency of CRISPR-mediated duplication compared to CRISPR-induced deletion^22,23,42^. We estimated deletion efficiencies for individual gRNAs RGD locus combining data from all relevant gRNAs (**Supplementary Tables 6-8**). These analyses demonstrate that the PB system can deliver multiple RGD cellular models in parallel, although the efficiencies of generating very large RGD rearrangements remain low, consistent with our initial publication of the SCORE method^22^.

**Figure 4.**
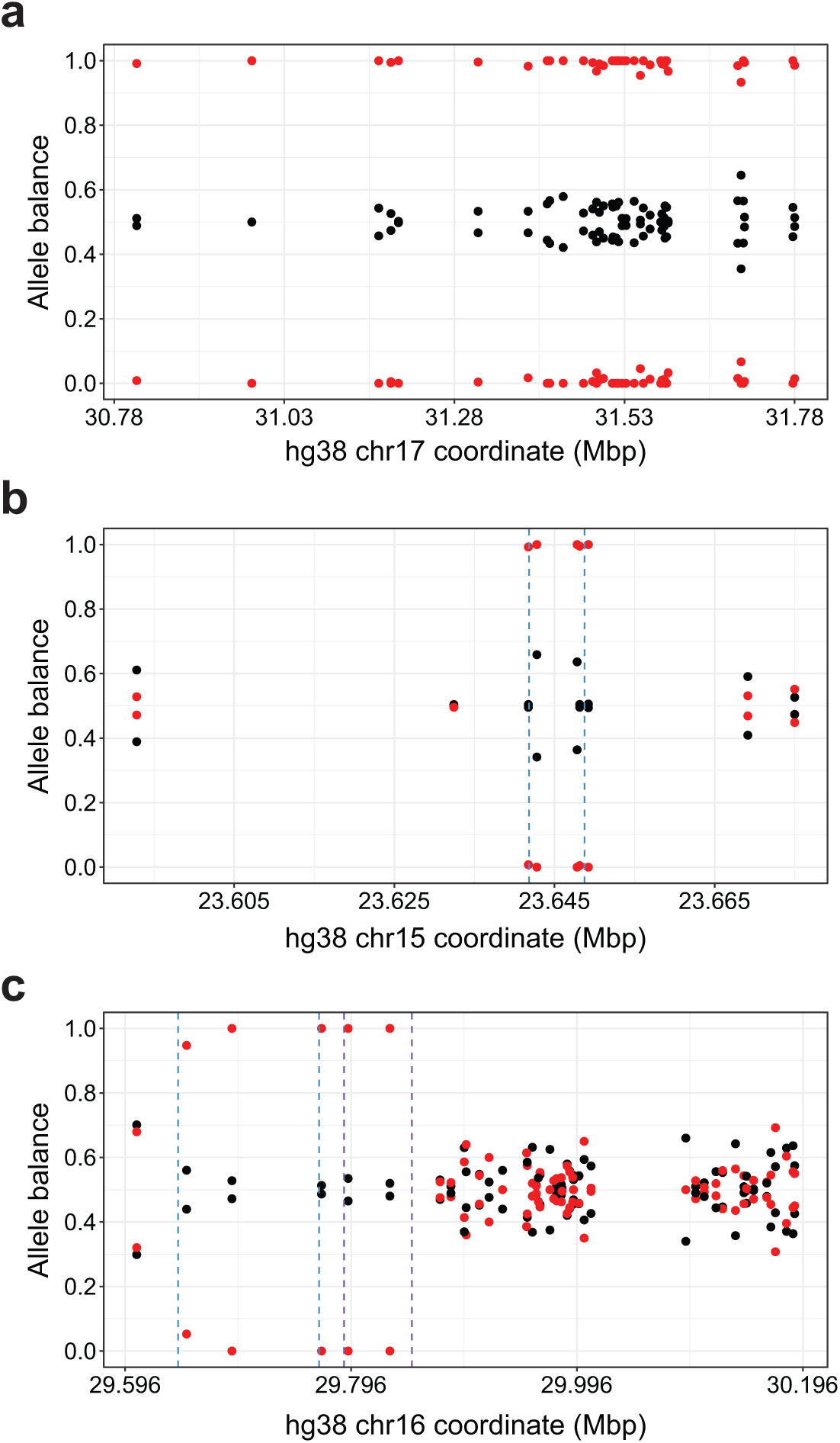
Selected PB CRISPR deletions. Plots depict allele balance (points) at MIP targets across genomic sequence for lines harboring a Cas9-induced deletion (red) and representative unedited controls (black). Dashed vertical lines indicate locations of gRNA cleavage sites. a) An ∼1.4 Mbp RGD deletion at chromosome 17q11.2 mediated by a single gRNA targeting flanking duplicated sequences (not shown). b) An ∼6.9 kbp deletion encompassing *MAGEL2* resulting from editing with a corresponding gRNA pair (blue). c) An ∼214 kbp deletion at chromosome 16p11.2 involving Cas9 activity directed by two separate gRNA constructs (blue, purple).

## DISCUSSION

We developed a method for establishing iPSC mutational models in parallel and engineered at least 56 lines with indels and 8 lines with genomic deletions in a single experiment. Compared to model-making via serial or arrayed gRNA transfections, our PB platform drastically reduces labor requirements and associated costs, consolidating cell culture experiments that could take many months. PB editing also obviates the need for repeated collection and storage of unedited control lines corresponding to each mutant, provided that obtaining mutant-control pairs exposed to the same gRNAs is not critical. Our strategy generated indels efficiently (31.1%) and deletions at lower rates (1.5% for RGD deletions and < 1% for focal deletions). These data are consistent with mechanistic considerations and previous work showing higher editing rates for indels than CNVs^43,44^, with RGD deletions made using a single gRNA produced more efficiently than focal deletions mediated by two gRNAs^22^. Importantly, we observed primarily single-copy indel formation, suggesting that we could efficiently obtain heterozygous loss of function mutants, which often model human disease more precisely than homozygous knockouts. Furthermore, we recovered 8 lines with indels at multiple loci (**Fig. 3b**), demonstrating that the PB system can deliver oligogenic models. By isolating more lines and/or performing more comprehensive genotyping, PB editing can easily be scaled to accommodate projects aiming to create as many as hundreds of mutant cellular models and unedited controls in a two-month timeframe.

We envision that PB genome engineering could be used to genetically dissect pathogenic CNV regions^45^ or disrupt sets of genes involved in a common biological pathway or statistically associated with a disease phenotype^3^. For example, the chromosome 16p11.2 RGD locus contains 25 protein-coding genes^6^. Inactivating each one via frameshifting indels would entail designing a library of 25 gRNAs, or 50 if including two gRNAs per gene to control for potential off-target effects. With one or two additional gRNAs targeting segmental duplications flanking the genes, this library would also mediate chromosome 16p11.2 RGD deletions^22^. Our simulations^46^ (**Supplementary Fig. 5** and **Supplementary Table 9**) indicate that for a library containing 52 gRNA constructs, 1,963 PB integrations on average would be necessary to achieve at least 25 integrations of each construct. Since we observed an average of 2.17 integrations per line, we estimate 905 lines would collectively harbor the desired number of integrations.

Refinements to our protocol may facilitate genomic dosage alteration, including modifying transfection conditions to attain more PB integrations^35,41^, enhancing gRNA expression levels, harnessing other CRISPR systems^47,48^, and/or lengthening the duration of editing by prolonging *Cas9* induction^49^. Although we found no relationship between PB DNA input and numbers of integrations per cell, multiple experiments exploring a wider range of PB input levels and alternative delivery techniques have reported higher per-cell PB integration counts^34-36^. Thus, PB integrations could be increased to provide more mutational opportunities for any fixed number of iPSC lines. Another report suggests that enhancing gRNA expression may promote more efficient CNV formation. Using a PB platform akin to that presented here to deliver single gRNA pairs to stem cells and induce editing yielded small deletions (< 3 kbp) at estimated rates above 40%^35^. Based on these data, having multiple integrated copies of a given gRNA pair per cell and resulting higher gRNA expression levels appear conducive to efficiently inducing the deletion of interest. Accordingly, approaches for boosting gRNA expression warrant further exploration, including incorporating multiple gRNAs into the same transcript^50^ and limiting CRISPR library size to probabilistically obtain cells with two or more integrations of specific gRNA constructs (**Supplementary Table 10**). Most notably, two recent studies demonstrated engineering of genomic deletions in human stem cells using type I-E CRISPR/Cas systems, with efficiencies up to 14.6% and sizes up to ∼100 kbp^47,48^. Combining this technology with PB delivery of corresponding gRNA libraries offers tremendous promise for further scaling deletion modeling in the future.

Deploying PB CRISPR libraries for parallelized production of cellular mutational models unlocks several exciting new possibilities. First, this method could be leveraged to generate compendiums of near-isogenic iPSC lines with deletions or frameshifting indels across hundreds of loci to investigate consequences of partial or complete loss of function. Second, library design and transfection conditions could be modified to promote frequent multiplex mutagenesis within cells, rendering oligogenic modeling more tractable and scalable. Third, with further development, we anticipate that our approach will accommodate base editing^20,21^ and prime editing^25^, expediting the creation of cellular models featuring patient-specific point mutations. Given the relatively high efficiency of these CRISPR strategies, they should dovetail well with our PB system. Fourth, PB libraries increase multiplexing capacity for CRISPR activation/interference applications^36,37^. In all these ways, our PB method will promote systematic efforts to compile large collections of cellular models in isogenic backgrounds. Such allelic series will serve as powerful resources for comparative functional genomics^51^, driving research into the impacts of genes and variants at unprecedented scope and scale.

## ACKNOWLEDGEMENTS

We thank T. Aneichyk, P. Boone, R. Collins, A. Domingo, S. Erdin, and C. de Esch for thoughtful discussion. This work was supported by U.S. National Institute of Health grants R01NS093200 (to J.F.G.) and R01HD096326 (to M.E.T) and a grant from the Foundation for Prader-Willi Research (to M.E.T.). X.N. was supported by a U.S. National Institute of Health fellowship F32MH115614. M.M.O. is supported by an Autism Speaks postdoctoral fellowship 11815. K.M. is supported by a U.S. National Science Foundation Graduate Research Fellowship DGE-1745303. R.Y. is supported by the Massachusetts General Hospital Fund for Medical Discovery.

## AUTHOR CONTRIBUTIONS

X.N. and M.E.T. designed the study. X.N. performed cloning experiments, including construction of the PB vector and assembly of gRNA libraries, with assistance from N.D.B. X.N. designed gRNAs and gRNA pairs. N.D.B. performed iPSC experiments, with assistance from X.N. M.M.O., K.M., and D.J.C.T. performed iPSC transfection experiments exploring a wide range of conditions and assisted with troubleshooting during early stages of method development. B.C. performed library construction for whole-genome sequencing (WGS) of the unedited iPSC line used in experiments. R.Y. performed SNV and indel calling from WGS data. X.N. designed MIPs, performed genotyping experiments, analyzed data, and performed simulations. X.N., N.D.B., J.F.G., and M.E.T. wrote the paper, with input and approval from all coauthors.

## COMPETING FINANCIAL INTERESTS

The authors declare no competing financial interests.

## METHODS

### Plasmid construction

We constructed the PB transposon plasmid pXN1Ds (Addgene deposition forthcoming) through a series of cloning steps modifying pPB-rtTA-hCas9-puro-PB (graciously provided by Dr. William Pu)^34^. Briefly, a gRNA cloning site derived from pPB-US-ECasE (a gift from Eleanor Chen, Addgene plasmid #83961) was inserted before the *TRE3G* promoter; sequence encoding an SV40 nuclear localization signal (NLS) was added to the beginning of the *Cas9* gene; internal *Cas9* genic sequence was replaced with corresponding sequence from PX459 (a gift from Feng Zhang, Addgene plasmid #48139) to eliminate BbsI restriction sites; sequence encoding the SV40 NLS at the end of the *Cas9* gene was replaced with sequence encoding a nucleoplasmin NLS; and the gRNA cassettes after *Cas9* were removed. We created the PB transposase plasmid pSPBase by removing the *ccdB* gene from PB210PA-1 (System Biosciences) to render it generally propagable. Finally, we developed the excision-only PB transposase plasmid pXPBase-GFP through multiple cloning steps modifying PB220PA-1 (System Biosciences). The *ccdB* gene was removed to render the plasmid generally propagable, sequence encoding a FLAG-tag at the transposase N-terminus was eliminated, and sequence encoding a T2A self-cleaving peptide and enhanced GFP was appended to the excision-only PB transposase gene. All plasmid constructs were validated through massively parallel sequencing performed by the MGH DNA core.

### gRNA library design

We designed two gRNA libraries: a library of single gRNAs to engineer microdeletions and microduplications of chromosomal regions associated with RGDs and a library of gRNA pairs to engineer deletions and duplications of single genes or sets of genes within a few of these regions (**Supplementary Table 1**). To design the first library, sequences of segmental duplications flanking these regions (and in some cases, also related paralogous sequences) were aligned using Clustal 2.1^52^ or MAFFT v7.310^53^. These alignments were parsed to identify all candidate gRNA sequences, defined as 20 bp sequences identical between target segmental duplications that precede a canonical 3’ NGG S. pyogenes protospacer adjacent motif (PAM) in both and lack an identical protospacer in other aligned paralogous sequences. gRNA candidates were then scored for predicted efficiency using Azimuth 2.0^54^ and initially assessed for potential off-target activity by concatenating protospacers with each possible NGG PAM and mapping them to the human reference genome GRCh38/hg38 using mrFAST^55^. All guide candidates having more perfect matches than a specified threshold were eliminated. To more carefully analyze possible off-target sites, we used Cas-OFFinder^56^ to identify all loci in GRCh38/hg38 with up to five mismatches to each candidate protospacer with its PAM, not counting mismatches at the degenerate first PAM position. Cas-OFFinder output was parsed to compute genome-wide counts of sequences having 0-5 mismatches to each candidate as well as the maximum Cutting Frequency Determination (CFD) score^54^ considering all identified off-target sequences for each candidate. We manually selected guides from our filtered set of candidates considering both their predicted activities (with higher Azimuth 2.0 scores better) and their likelihoods of specific targeting (with lower maximum CFD scores and fewer off-target sites with 1-2 mismatches better). gRNA design for the second library (gRNA pairs) employed a similar framework, except candidates were initially derived from single sequences rather than sequence alignments, candidates within genes were excluded, and lower thresholds were applied to eliminate candidates based on perfect matches identified using mrFAST mapping.

### gRNA library cloning

DNA oligonucleotides (oligos) corresponding to single gRNAs (**Supplementary Table 1**) were ordered (Integrated DNA Technologies), pooled, and amplified using PCR primers Oligo-Fwd-2 (5’-GGCTTTATATATCTTGTGGAAAGGACGAAACACC-3’) and Oligo-Rev-2 (5’-GCCTTATTTTAACTTGCTATTTCTAGCTCTAAAAC-3’). pXN1Ds was digested with BbsI-HF (New England BioLabs), followed by gel purification of the vector backbone. We cloned our single-guide amplicon library into this vector using NEBuilder HiFi DNA Assembly Master Mix (New England BioLabs), incubating the Gibson assembly reaction at 50°C for 30 minutes. The resulting plasmid library was transformed into NEB Stable Competent *E. coli* (New England BioLabs), and 1% of the transformation reaction was plated to estimate library coverage by counting colonies. The remainder was used to inoculate a 100 mL culture (LB with 100 µg/mL carbenicillin) grown in a shaking incubator (250 rpm) at 30°C for 20-24 hours. Endotoxin-free plasmid library DNA was prepared using the ZymoPURE II Plasmid Maxiprep Kit (Zymo Research). To clone the library of gRNA pairs, we adapted previously published cloning workflows^43,57^. Briefly, we ordered DNA oligos (Integrated DNA Technologies) corresponding to each gRNA (**Supplementary Table 1**) as well as a gBlock (Integrated DNA Technologies) containing gRNA backbone and H1 promoter sequence. Each oligo pair was ordered in a separate well, and all oligos shared a common 18 bp of overlap with their partners at their 3’ ends. Oligo pairs were converted to double-stranded fragments containing linked gRNA pairs separated by an Esp3I cloning site via a single cycle of PCR. These fragments were then pooled, amplified, and cloned into an intermediate vector using Gibson assembly with the reaction incubated for 1 hour. Next, the resulting plasmid library and gBlock were digested with Esp3I (New England BioLabs) and gel purified. We cloned the gBlock insert into our vector library using the Quick Ligation Kit (New England BioLabs) and transformed and prepared DNA from this plasmid library as detailed above. Finally, we amplified final gRNA pair constructs using PCR, cloned them into pXN1Ds using Gibson assembly, and transformed and prepared DNA from this final gRNA pair plasmid library as detailed above. Colony counts indicated both single-guide and dual-guide plasmid libraries comfortably exceeded 20x library coverage.

### iPSC culture

iPSCs were cultured on plates (Falcon) coated with Matrigel hESC-Qualified Matrix (Corning) in Essential 8 medium (E8, Gibco) supplemented with penicillin/streptomycin (P/S, Biological Industries). For routine culture, cells were maintained on 6-well plates in a humidified incubator at 37°C with 5% CO2 and passaged every 3-4 days (at ∼80% confluence) using ReLeSR (Stemcell Technologies), following the manufacturer’s protocol. Working medium (E8 + P/S) was refreshed daily. Upon thawing and for the first 2-4 passages thereafter, cells were plated into working medium with 10 µM ROCK inhibitor (RI, Y-27632, Biological Industries); working medium was used for all other routine passages.

### gRNA library transfection

For each transfection, we seeded iPSCs into a new well in a 6-well plate to achieve ∼20% confluence upon plating – performing ReLeSR treatment, breaking colonies into 5-10-cell clumps, and plating into working medium with 10 µM RI. Twenty-four hours later, cells were transfected with PB plasmids using Lipofectamine Stem reagent (Invitrogen) according to the manufacturer’s protocol. We performed six transfections: (1) transposase-only control, 1000 ng pSPBase; (2-5) experimental series, 1000 ng pSPBase with 125 ng, 250 ng, 500 ng, or 1000 ng of PB gRNA plasmid libraries; and (6) transposon-only control, 1000 ng PB gRNA plasmid libraries. All transfections with PB gRNA plasmid libraries included equal nanogram amounts of single-guide and gRNA pair libraries. Cells were exposed to transfection complexes for 24 hours.

### Puromycin selection and genome engineering

Three days after PB plasmid transfections, for each condition, we performed single-cell passaging using accutase (Biological Industries) to seed 750,000 cells into a new well in a 6-well plate. Twenty-four hours later, each well was treated with working medium containing puromycin (ThermoFisher) at a concentration of 0.5 µg/mL. Working medium with puromycin was refreshed daily until no live cells remained in control wells (∼3-4 days of puromycin treatment total). Each well of puromycin-resistant iPSCs was grown for four days and then passaged following our ReLeSR seeding protocol above, except plating medium also contained doxycycline hydrochloride (Sigma-Aldrich) at a concentration of 2 µg/mL. Doxycycline-containing medium was refreshed each of the next two days, such that cells underwent doxycycline treatment and *Cas9* induction for three days.

### FACS and 96-well iPSC culture

After doxycycline treatment, colonies were dissociated into single cells using accutase, resuspended in Dulbecco’s Phosphate Buffered Saline (DPBS, Gibco) with 10 µM RI, and stained with TO-PRO-3 viability dye (ThermoFisher). Cells were then filtered through a 35 µm cell strainer (Corning) and sorted using a BD FACSAria II Flow Cytometer (BD Biosciences) equipped with a 100 µm nozzle. We sorted at 20 psi under sterile conditions for live (TO-PRO-3-), single iPSCs, setting the instrument to deposit one cell into each well in a 96-well plate containing working medium with 10% CloneR cloning supplement (Stemcell Technologies). Filled plates were transferred to the 37°C incubator and left undisturbed for 48 hours after sorting. Medium with CloneR was refreshed daily on days 2-3 post-FACS, and then starting on day 4, medium changes (without CloneR) were performed every other day. On day 14, we passaged visible colonies using ReLeSR to consolidate them into new 96-well plates. For these passages, cells were plated into working medium with 10 µM RI.

### 96-well DNA extraction

DNA extraction from clonal iPSC lines cultured in 96-well plates was performed using the Quick-DNA 96 Kit (Zymo Research). Briefly, we treated colonies with ReLeSR to detach them from the plate and resuspended them in genomic lysis buffer. Subsequent steps involving cell lysis, DNA isolation, washing, and elution were performed following the manufacturer’s protocol.

### WGS

WGS was performed using a modified NEBNext UltraII for DNA Library Prep kit (E7645L, New England BioLabs). Briefly, 1 µg of DNA from the unedited iPSC line was fragmented to approximately 350 bp using Covaris shearing (E220, Covaris). DNA fragments were end repaired and A-tailed using Ultra II reagents. Illumina stubby-Y adapters were ligated to fragments and finished with PCR using 8-bp barcoded primers (10005921, Integrated DNA Technologies) using Ultra II reagents. Individual libraries were pooled and sequenced on NovaSeq S4 using pair-end 150 bp chemistry on the Walk-Up Sequencing Platform at the Broad Institute.

### SNV calling

WGS data were quality controlled using FastQC^58^. Read pairs obtained from sequencing were aligned to GRCh37v71 with BWA-MEM v0.7.10-r789^59^. Alignments were processed using PicardTools^60^ and samtools^61^ to check the quality of the alignments. The Genome Analysis Toolkit (GATK)^62^ v3.5 was applied for base quality score recalibration, indel realignment, duplicate removal, and SNV and indel discovery and genotyping as per published best practice protocols. GATK-called variants were filtered based on base quality, sequencing read depth, genotype quality, results of a rank-sum test for relative positioning of reference (REF) versus alternate (ALT) alleles within reads, strand bias, and allelic bias.

### MIP genotyping

We designed 2,022 single-molecule MIPs^40^ (**Supplementary Table 11**) to simultaneously genotype sequence at several Cas9 target sites, copy number across genes and genomic regions targeted for dosage alteration^39^, the presence/absence of PB transposon integrations, and identities of integrated gRNA sequences. A few MIPs also targeted pXN1Ds sequences outside the PB transposon to enable detection of any random plasmid integration if it occurred. All MIPs were designed as previously described^39^, except MIPs assaying copy number across unique genomic sequence were designed to target SNVs present in one or both iPSC lines widely used in our lab. MIP pooling, capture, and sequencing were performed as previously described^39^ using DNA extracted from clonal iPSC lines cultured in 96-well format as input.

### MIP analyses

We mapped MIP sequencing data to a custom genome containing four mock chromosomes per unique chromosomal region of interest, with each mock chromosome having one of the four DNA nucleotides at all SNV positions. This custom genome also incorporated multiple duplicated sequences successfully genotyped in other samples using MIPs to serve as positive controls and provide quality control for each sample. Lastly, this custom genome included all gRNA target sites, all pXN1Ds sequence, and mock chromosomes corresponding to each gRNA construct. To determine which gRNA constructs were integrated in each iPSC line, we counted numbers of capture events corresponding to each gRNA expressed from a U6 promoter and imposed minimum count thresholds for calling each gRNA construct as integrated. Using a similar strategy, we assessed each line for random plasmid integrations by counting capture events corresponding to pXN1Ds sequences within and outside the PB transposon (four MIP targets each). Targets exceeding a minimum count threshold were called as genomically present. If all four targets within the transposon were present and all four targets outside the transposon were absent, the integration was called as a PB integration. Otherwise, we deemed the iPSC line as harboring no integrations (all eight targets absent) or containing random plasmid integration (all other scenarios). Finally, we inferred CNVs from altered SNV allele balance over targeted genomic intervals and characterized Cas9-induced indels by sequence analysis at Cas9 cleavage sites. To prevent misinterpretation of indels arising from sequencing errors, all mapping alignments below specified capture event count and allele fraction thresholds were discarded prior to indel calling.

### PB experiment simulations

We implemented a Markov chain^46^ to simulate PB experiments with different gRNA library sizes and different desired integration profiles. Briefly, our model randomly integrates one gRNA construct at a time until each construct in the library reaches a minimum number of integrations. Running this simulation 100,000 times for each experimental scenario and averaging the numbers of integrations required yielded estimates of how many integrations are necessary to achieve prespecified outcomes pertaining to gRNA library integration. Thus, these simulated data are useful for designing new PB gRNA libraries and planning corresponding experiments, e.g., determining how many iPSC lines to isolate.

---

**Supplementary Figure 1.**
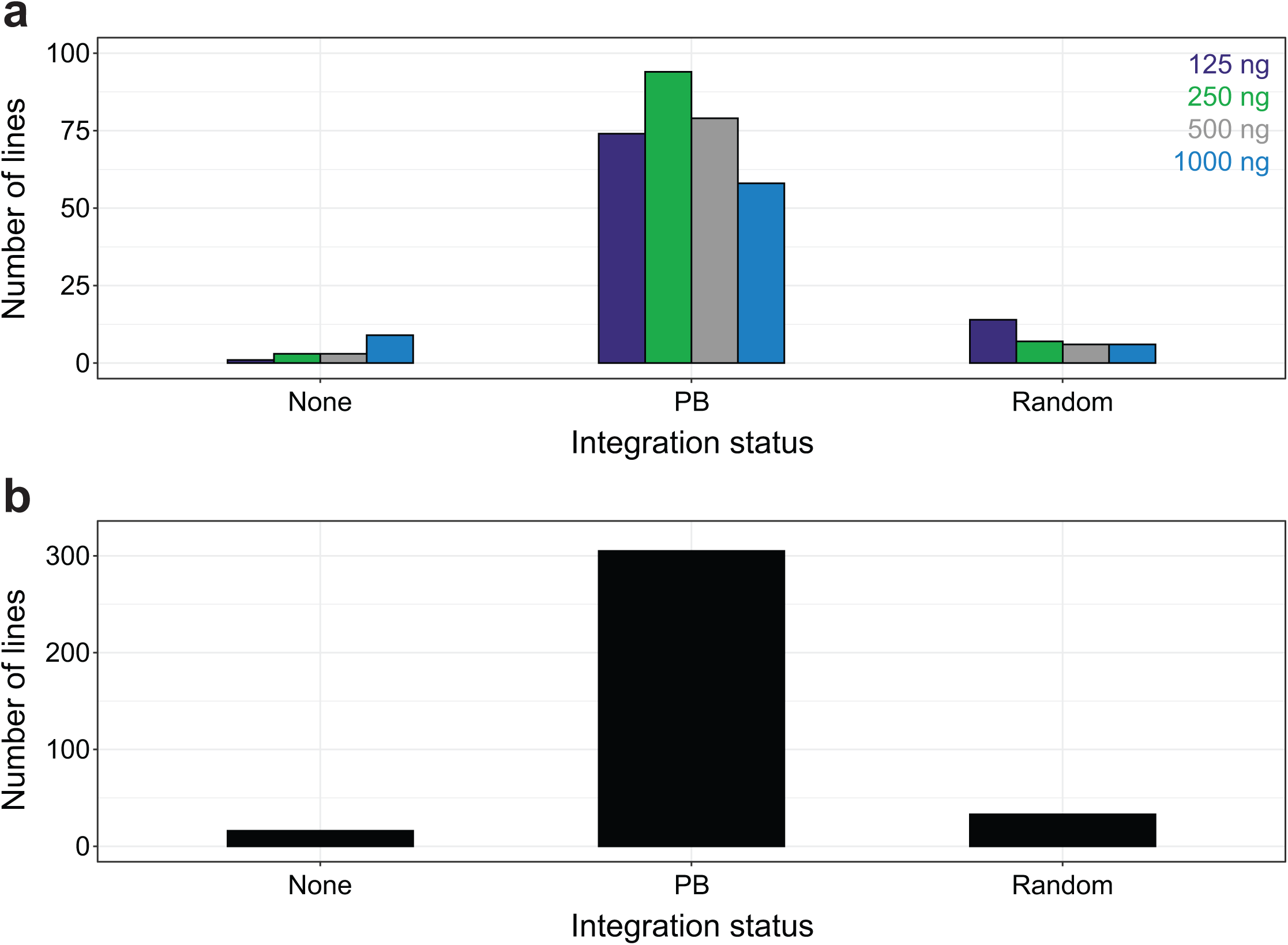
Integrity of PB integrations. iPSC lines were classified as lacking PB integrations (none), containing PB integrations arising from transposition (PB), or harboring random plasmid integrations (random) based on the genomic presence or absence of PB vector sequences within and outside the PB transposon. Distributions illustrate numbers of lines assigned to each category a) stratified by transfection conditions and their associated PB DNA input levels and b) aggregated across all transfections.

**Supplementary Figure 2.**
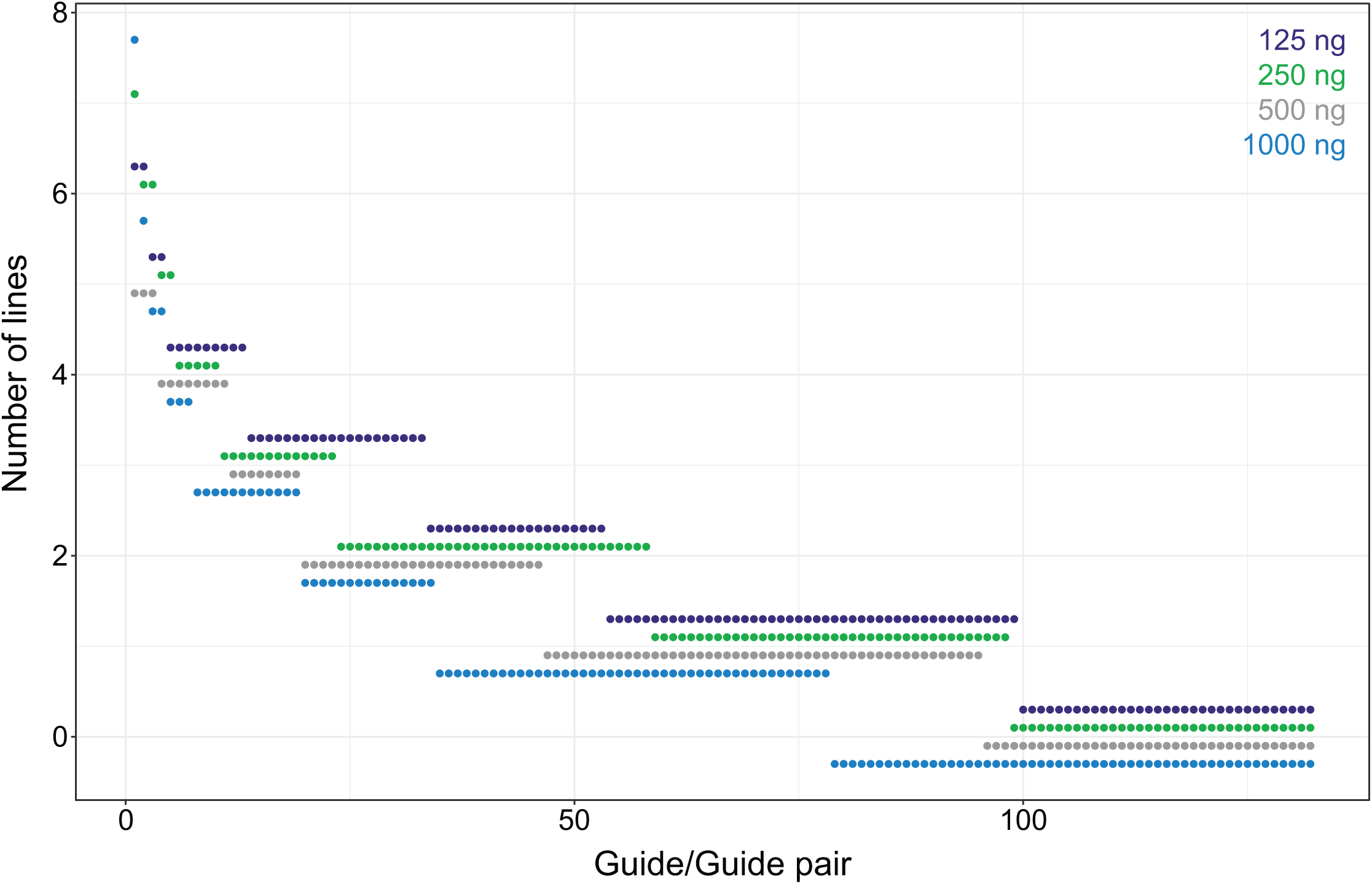
gRNA library representation for each PB input level. Points show numbers of lines derived from each transfection containing each gRNA construct. Constructs are ranked separately for each condition based on abundance.

**Supplementary Figure 3.**
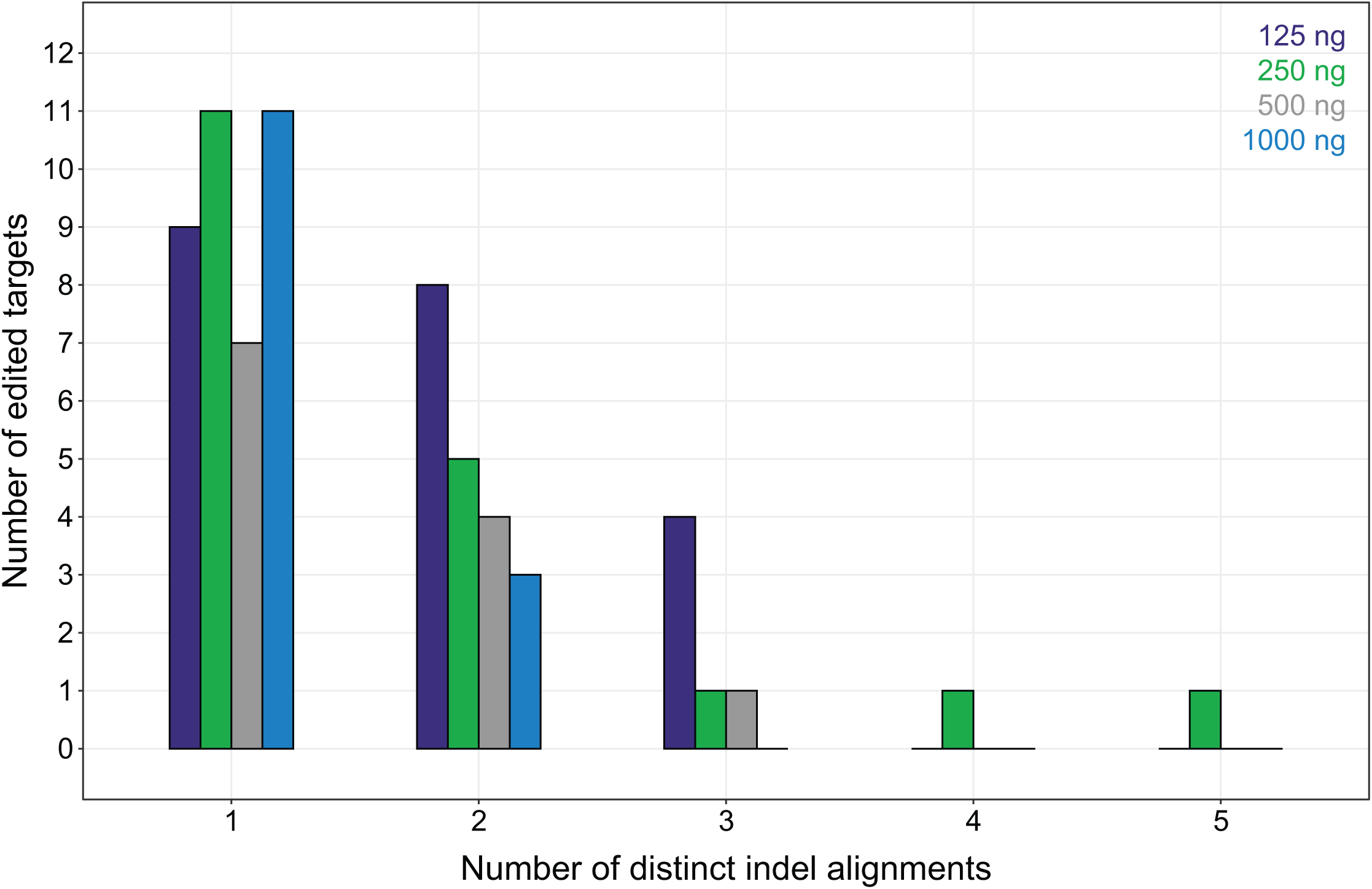
Breakdown of indel editing. Distribution of numbers of distinct indel alignments per edited target per iPSC line. Data are stratified based on PB DNA input levels used for transfection. Multiple indel alignments most frequently reflect distinct indels arising from editing activity at different copies of the same target site.

**Supplementary Figure 4.**
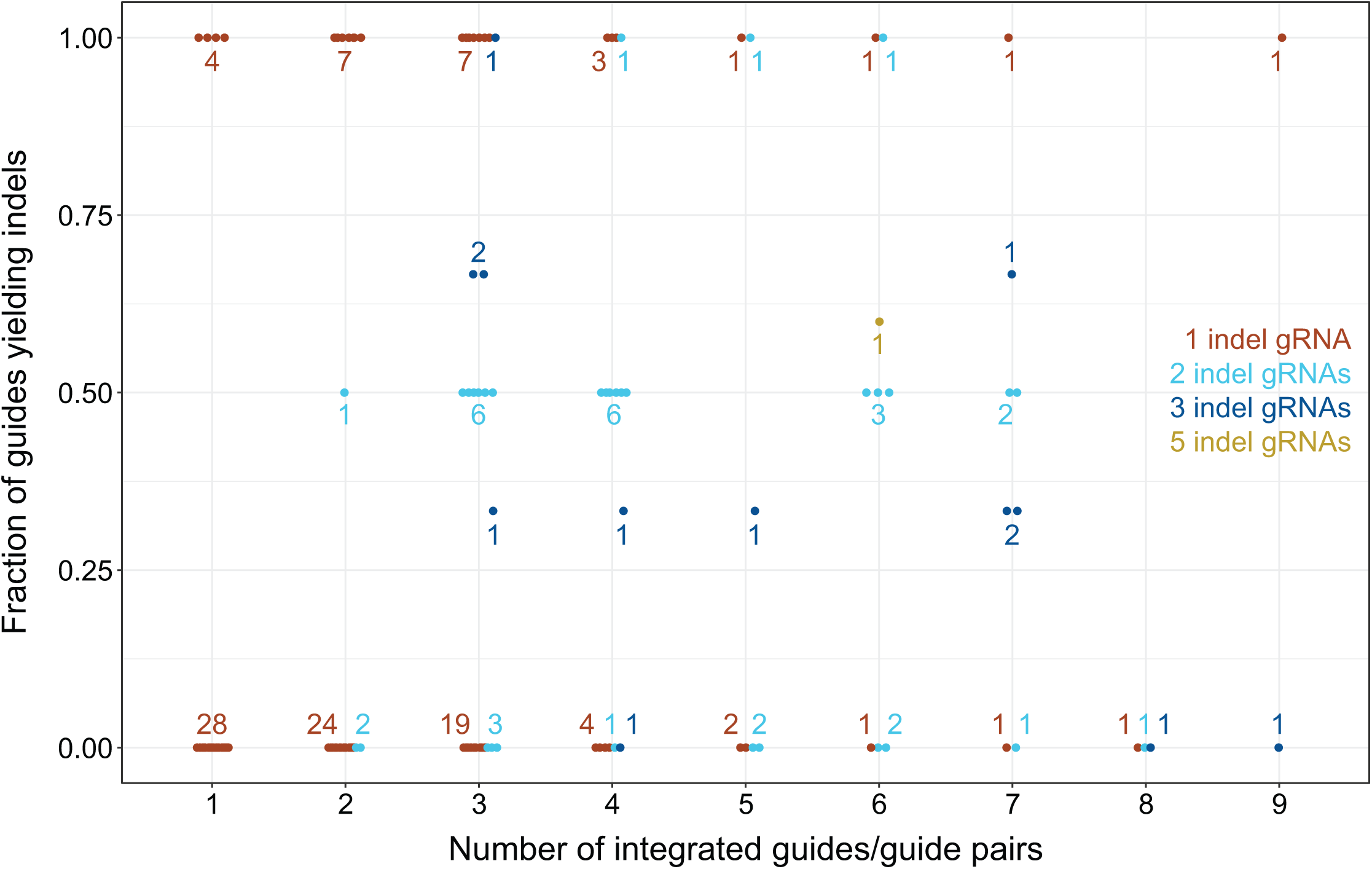
Indel editing efficiencies in lines with different PB copy numbers. Each point denotes a single iPSC line expressing one (orange) or more (cyan, blue, gold) gRNAs amenable to indel analysis (indel gRNAs). Positions indicate fraction of the indel gRNAs in that line that mediated indel formation (y axis) versus the total numbers of gRNA constructs expressed in that line (x axis, data jittered around their integer values). Colored numbers quantify corresponding points at each position.

**Supplementary Figure 5.**
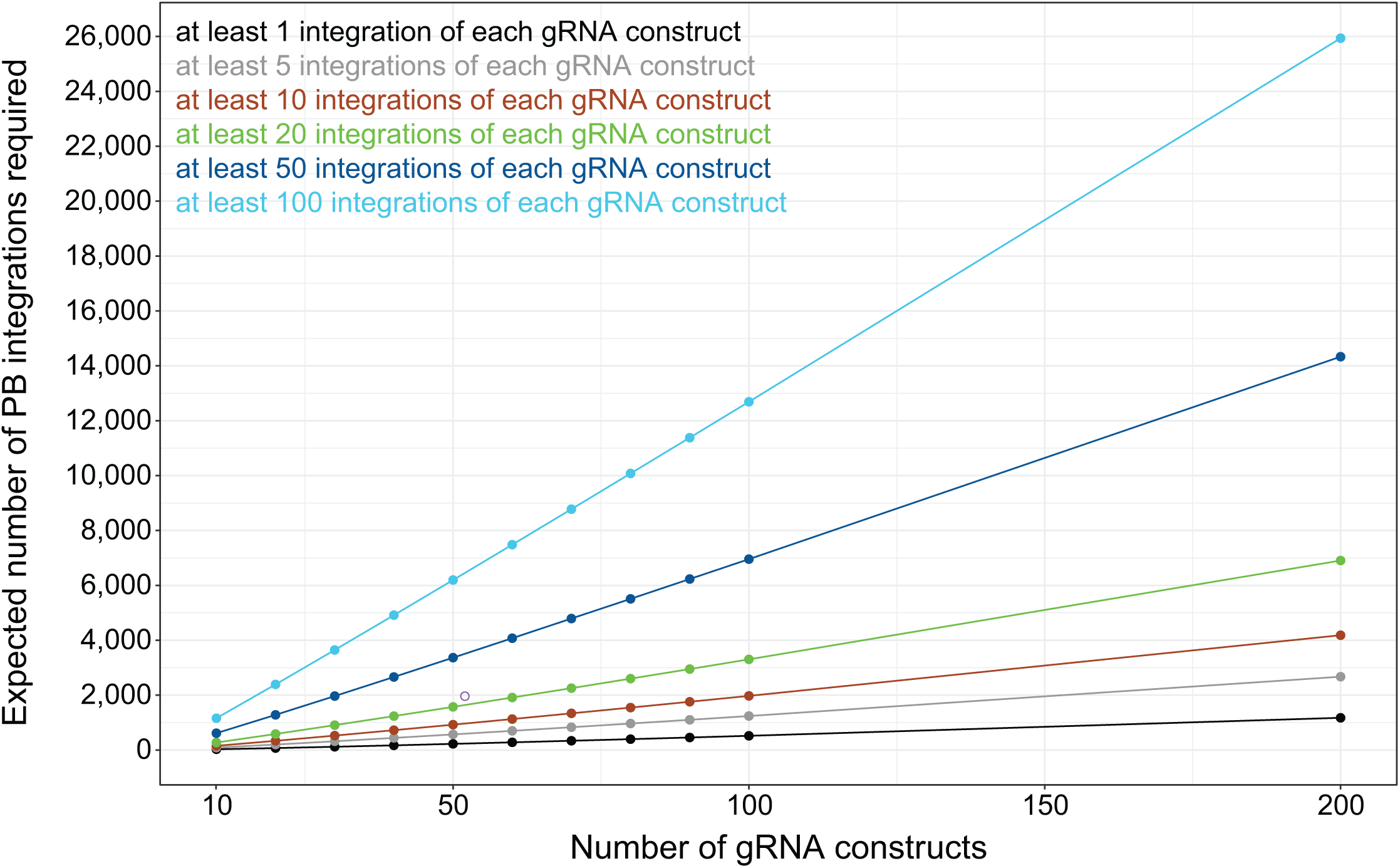
Estimates of desired numbers of PB integrations under various experimental scenarios. Each point represents the average (from 100,000 simulations) number of PB integrations needed (y axis) to achieve a minimum specified amount of integrations (color) of each gRNA construct, considering different gRNA library sizes (x axis). Open purple circle corresponds to chromosome 16p11.2 gRNA library example presented in the discussion.

## Notes

### Competing Interest Statement

The authors have declared no competing interest.

